# Differential effects of two HDAC inhibitors with distinct concomitant DNA hypermethylation or hypomethylation in breast cancer cells

**DOI:** 10.1101/578062

**Authors:** Arunasree M. Kalle, Zhibin Wang

## Abstract

DNA methylation and histone acetylation are the two important epigenetic phenomena that control the status of X-chromosome inactivation (XCI), a process of dosage compensation in mammals resulting in active X chromosome (Xa) and inactive X chromosome (Xi) in females. While DNA methyltransferases (DNMTs) are known to maintain the DNA hypermethylation of Xi, it remains to be determined how one or a few of 18 known histone deacetylases (HDACs) contribute(s) to Xi maintenance. Herein we found that HDAC1/2/4/6 were overexpressed in breast cancer cells, MDA-MB-231, with Xa/Xa status compared to normal breast epithelial cells, MCF10A, with Xa/Xi status. Inhibition of these overexpressed HDACs with two different drugs, sodium butyrate (SB) and Trichostatin A (TSA), caused surprisingly distinct effects on global DNA methylation: hypermethylation and hypomethylation, respectively, as well as distinct effects on a repressing histone mark H3K27me3 for heterochromatin and an active mark H3K56ac for DNA damage. Surveying three DNMTs through immunoblot analyses for insights revealed the up- or down-regulation of DNMT3A upon drug treatments in a concentration-dependent manner. These results correlated with the decreased *XIST* and increased *TSIX* expression in MDA-MB 231 as a possible mechanism of Xi loss and were reversed with SB treatment. Further RNA-seq analysis indicated differential gene expression correlating with the promoter methylation status of a few genes. Collectively, our results demonstrate a crosstalk between HDACs and DNMTs and the novel involvement of HDACs in skewed Xi in breast cancer.

## Introduction

Lyonization is the phenomenon of X chromosome inactivation (XCI) in female mammals to compensate the dosage of genes on X chromosome in both the genders [1]. During early embryogenesis, the initiation of XCI occurs by counting the number of X chromosomes followed by random selection of an X chromosome to be inactivated (Xi) [2]. Spreading of the long non-coding RNA XIST on Xi and inhibition of XIST on active X chromosome (Xa) by another long non-coding RNA TSIX establishes the inactivation [3]. Once Xi is established, it is ensured that the same X chromosome is inactivated in progeny cells after cellular division by DNA hypermethylation and histone hypoacetylation [4–6].

DNA methylation patterns, maintained by three DNA methyltransferases (DNMTs) [7–9], are also involved in Xi. Specifically, the *de novo* DNMT3A and DNMT3B are implicated in initiation of Xi whereas DNMT1 is involved primarily in maintenance of Xi chromosome [10, 11]. Of late, study has identified DNMT1 being as transcriptional activator of *XIST* gene [12] and therefore loss of DNMT1 results in Xi DNA hypomethylation.

Histone deacetylases (HDACs) are a family of proteins involved in deacetylation of histones on expressed and poised genes as demonstrated by us previously [13, 14]. There are 18 HDAC isoforms in humans classified into four classes: Class I (HDAC1, 2, 3, 8); Class II (HDAC4, 6, 7, 9, 10); Class III (Sirtuins, Sirt1 – Sirt7) and Class IV (HDAC11) [15]. Aberrant expression of these HDACs has been identified in many pathological diseases including cancer, diabetes, and neurodegenerative disorders [16]. Several studies have implicated the role of class I and class II HDACs in breast cancer. [17, 18]. Although all HDACs share a common function of gene silencing, the involvement of specific individual or group of HDAC isoforms in Xi maintenance remains to be determined.

Many studies have reported Xi reactivation [19], loss of *XIST* expression and hypomethylation of Xi DNA in breast cancer cells implicating loss of Xi maintenance [20]. *DNMTs* and *HDACs* are overexpressed in breast cancers, yet there is a loss of Xi. To the interplays of overexpressed *DNMTs* and *HDACs* and skewed Xi involved in breast cancer, herein we demonstrate a crosstalk between DNA methylation and histone acetylation *via* DNMT3A and HDAC1/ HDAC2 in chromatin remodeling complex, NuRD, in breast cancer cells. Inhibition of HDACs by sodium butyrate induces *de novo* methyltransferase DNMT3A by inhibiting UHRF1 ubiquitin ligase and thereby induces *XIST* expression in breast cancer cells.

## Results

### Differential expression of Class I and II HDACs in normal and cancerous breast cells

To understand the role of HDACs in Xi maintenance, we profiled class I and class II HDACs in normal and cancerous breast cells in the presence or absence of two different pan-HDAC inhibitors, sodium butyrate (SB) and Trichostatin A (TSA). Immunoblot analysis suggested that HDAC1 and HDAC2 of class I (Fig. 1A) and HDAC4 and HDAC6 of class II (Fig. 1B) HDACs were overexpressed in cancer cells when compared to normal MCF10A breast cells. There was no significant difference in other class I and class II HDACs.

**Fig 1:**
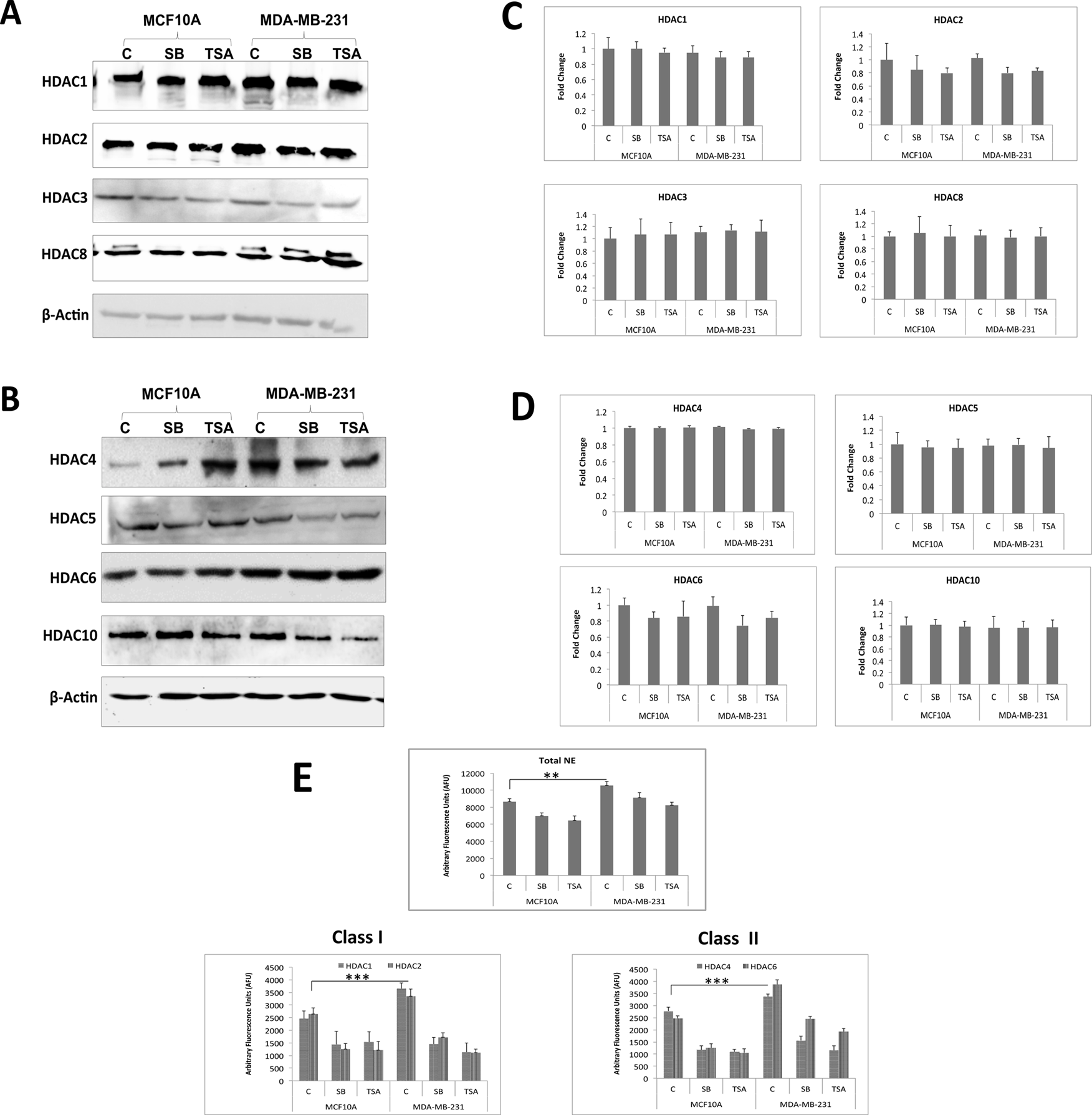
Increased expression and activity of class I and class II HDACs in normal and cancerous breast cells. **A.** Immunoblot analysis of class I HDACs. **B.** Protein expression levelss of Class II HDACs. **C.**Relative mRNA expression of class I HDACs. **D.** Quantitative mRNA expression analysis of class II HDACs. **E.** Graph showing the total HDAC activity in nuclear extracts of normal and cancers breast cells (Top). There was an increased class I (bottom left) and class II (bottom right) HDAC activity in breast cancer cells. C. Control, C,Sodium butyrate, SB, Trichostatin A,TSA.

To determine the extent to which different levels of HDAC1, HDAC2, HDAC4, and HDAC6 attributes to different RNA transcripts, we performed real-time PCR and results showed no notable differences of HDAC1, HDAC2, HDAC4 and HDAC6 mRNA among normal and cancer cells. However, SB and TSA resulted in reduction of HDAC2 and HDAC6 mRNA levels in both MCF10A and MDA-MB-231 cell lines, but not HDAC1 (Fig. 1C) and HDAC4 (Fig. 1D) No significant change in mRNA levels was observed in other class I and class II HDACs.

Next, we assayed the HDAC activity of total nuclear extracts along with activity of overexpressed HDAC1, HDAC2, HDAC4 and HDAC6 (immunoprecipitated from MCF10A and MDA-MB-231 cells. The activity assay results are in correlation with the real-time PCR and immunoblot analysis showing increased activity of the overexpressed HDACs in MDA-MB-231 cells compared to MCF10A cells (Fig. 1E).

Collectively, our RT-PCR, immunoblot and activity assay results suggest that the overexpression of HDAC1/2/4/6 in cancer cells is not due to increased transcription but probably due to increased protein stability and/or increased translation leading to increased deacetylation activity.

### Inhibition of HDACs by SB and TSA altered the expression of XIST (Xi), TSIX (Xa), BRCA1 and H3K56 acetylation

SB and TSA inhibited four (HDAC1, HDAC2, HDAC4 and HDAC6) out of eight examined HDACs. To decipher the role of these HDACs in Xi maintenance, we first analyzed the transcript levels of Xi (*XIST*) and Xa-specific genes (*TSIX*, *PGK1*, and *G6PD*) by real-time PCR. The results indicated that MDA-MB-231 cells lack *XIST* expression with considerable level of *TSIX* mRNA. On the other hand, MCF10A cells have observable levels of *XIST* and *TSIX* and their levels are similar. Next, we examined the effects of SB and TSA inhibition on the expression of Xi and Xa specific genes (Fig. 2). Significant increase in *XIST* and decrease in *TSIX* expression in MDA-MB-231 cells was observed when treated with SB. However TSA did not show much effect on the expression of these genes (Fig. 2A).

**Fig 2:**
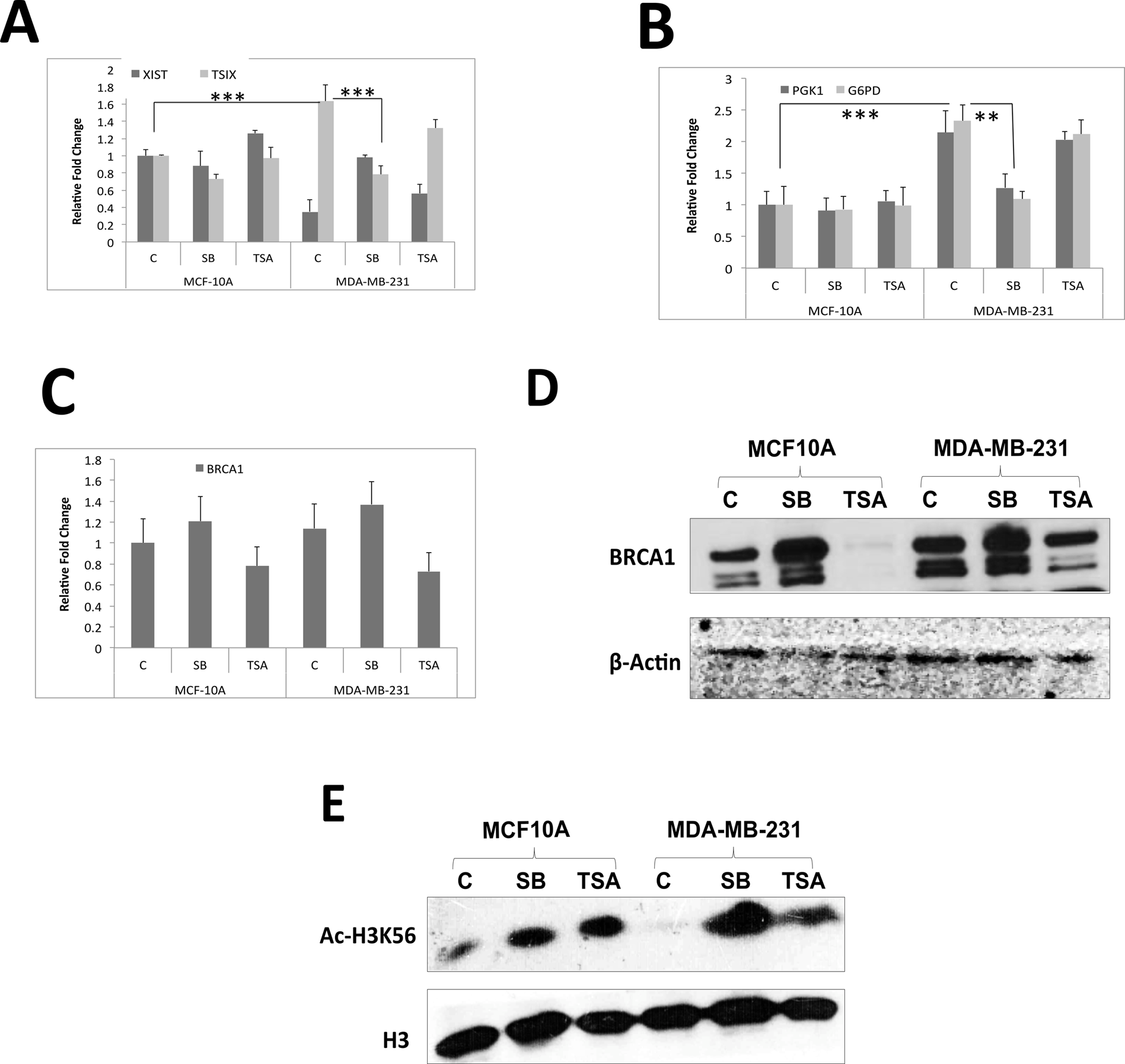
Effect of HDAC inhibition on Xi-/Xa-specific and BRCA1 genes. **A.** Effect of sodium butyrate (SB) and trichostatin A (TSA) on mRNA expression of XIST and TSIX long non-coding RNAs (lncRNAs) in cancer (MDA-MB-231) and normal (MCF10A) breast cells. Untreated cells served as control (C). **B.** Quantitative mRNA analysis of Xa-specific genes, G6PD and PGK1, in repsonse to Sodium butyrate (SB) and trichostatin a (TSA) in MCF10A, and MDA-MB-231 breast cells. **C.** Effect of Sodium butyrate (SB) and trichostatin a (TSA) on the expression of BRCA1 mRNA in MCF10A and MDA-MB-231 breast cells. C: Control untreated cells. **D.** Effect of Sodium butyrate (SB) and trichostatin a (TSA) on the expression of BRCA1 protein in MCF10A and MDA-MB-231 breast cells. C: Control untreated cells. **E.** Immunoblot showing Ac-H3K56 levels in MCF10A and MDA-MB-231 cells in response to Sodium butyrate (SB) and trichostatin A (TSA). Untreated cells served as control (C).

To further analyze the reversal of Xa to Xi by HDAC inhibition, we also evaluated the mRNA levels of Xa-specific genes *PGK1* and *G6PD*. Both *PGK1* and *G6PD* were upregulated in MDA-MB-231 cells indicating loss of Xi and duplication of Xa or reactivation of Xi. SB reduced the expression of *PGK1* and *G6PD* in MDA-MB-231 cells. But TSA did not show any significant reduction on *G6PD* and *PGK1* expression (Fig. 2B).

Breast Cancer 1 (BRCA1) is known to co-localize and transport *XIST* RNA onto Xi [21, 22] and loss of BRCA1 leads to Barr body (Xi) loss, high-grade tumor progression and chemoresistance [19, 23]. Studies have shown the loss of Xi and BRCA1 in basal-like breast tumors [24]. MCF10A and MDA-MB-231 are basal-like normal and cancerous breast epithelial cells, respectively. Therefore, we analyzed the expression of *BRCA1* at RNA and protein level in response to SB and TSA treatment. The transcript levels of *BRCA1* gene were not different in MCF10A and MDA-MB-231. TSA caused a decrease in BRCA1 RNA level in both the cell lines. However, SB caused a slight increase in *BRCA1* RNA levels (Fig. 2C). Intriguingly, BRCA1 protein levels were high in MDA-MB-231 compared to MCF10A cells. TSA caused a decrease in the BRCA1 protein levels significantly in MCF10A cells compared to MDA-MB-231 cells. Similar to the RNA levels, there was an increase in BRCA1 protein level in MDA-MB-231 and MCF10A cells treated with SB (Fig. 2D) and this result is in well agreement with increase in *XIST* expression (Fig. 2A).

The loss of BRCA1 and decreased H3K56 acetylation (H3K56ac) levels at DNA double strand breaks (DSB) promote Non-homologous-end joining (NHEJ), leading to genomic instability [25, 26]. To assesse the levels of H3K56ac in normal and cancerous breast cell lines, we did immunoblot with anti-H3K56ac in both cell lines. Data showed much lower level of H3K56ac in cancer cells when compared to MCF10A normal cells, implying genome instability in MDA-MB-231 cells. As expected, in response to SB and TSA treatment, both treatments increased the levels of H3K56ac significantly. Particularly, SB induced significantly when compared to TSA especially in MDA-MB-231 cells (Fig. 2E). These data suggest that SB treatment had a better effect on inhibiting NHEJ, thereby protecting genomic stability.

### HDAC inhibition resulted in unexpected, global DNA hypermethylation

Increased histone acetylations have been linked to the reversal of DNA hypermethylation [27]. Furthermore, inhibition of HDACs by SB and TSA showed an increase in *XIST* and decrease in *TSIX* expression along with reduction in *PGK1*, *G6PD* and *BRCA1* genes (Fig. 2). These observations prompted us to determine the effects of SB and TSA on DNA methylation in the two cell lines we used here. As expected, our dot blot analyses showed DNA hypomethylation in cancerous MDA-MB-231 breast cells compared to normal MCF10A breast cells (Fig. 3A). Consistent with early investigations [28], TSA treatment decreased global DNA methylation in both normal and cancerous breast cells (Fig. 3A). Intriguingly, significant increase in the global DNA methylation levels was observed in SB-treated cells.

**Fig 3:**
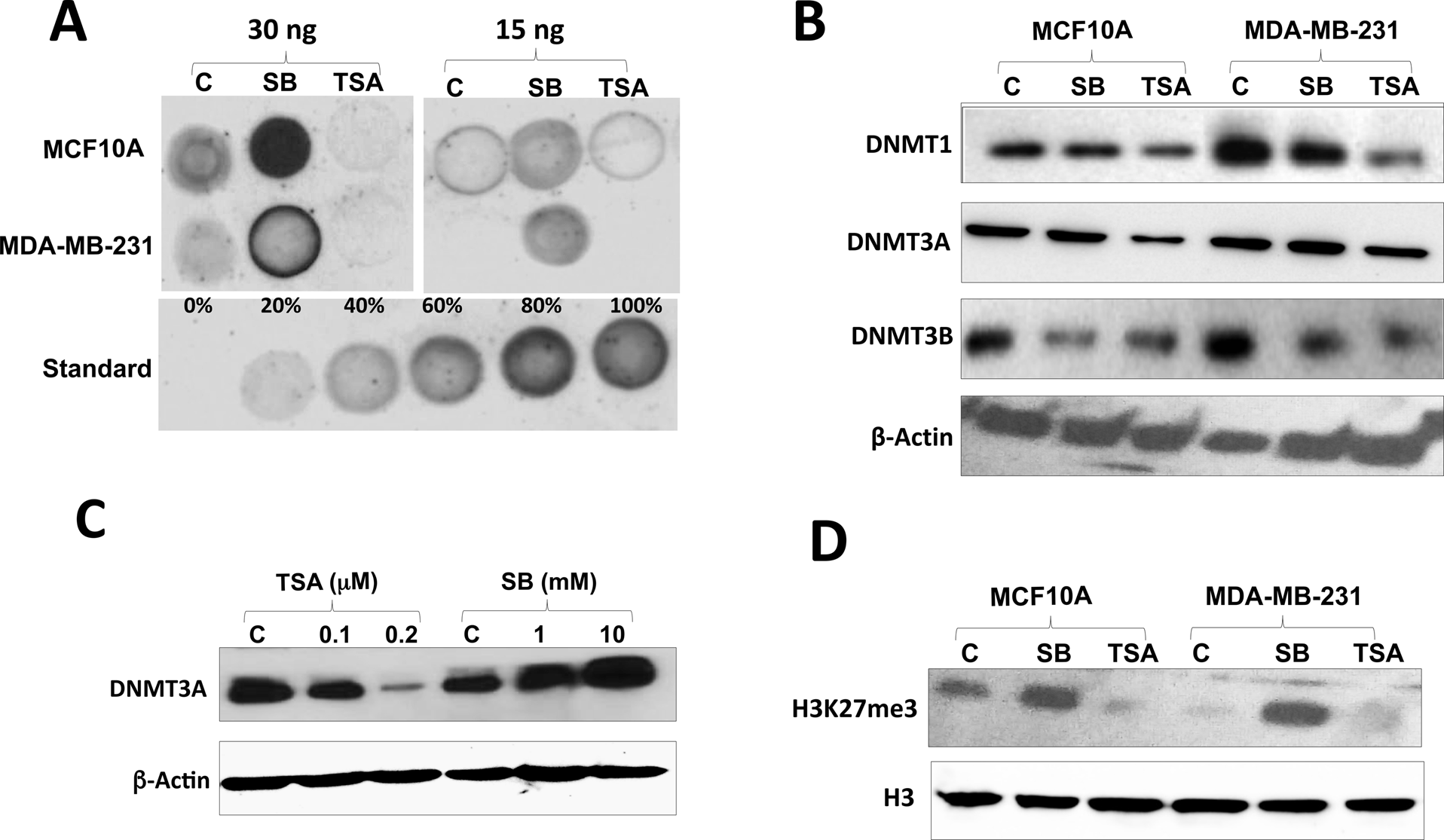
Differential effects of SB and TSA on DNA global methylation. **A.** DNA Dot blot showing the differential effects of Sodium butyrate (SB) and Trichostatin A (TSA) on global DNA methylation. Std: 100% methylated mouse genomic DNA. **B.** Effect of Sodium butyrate (SB) and trichostatin A (TSA) on the expression of DNMT proteins in MCF10A, MDA-MB-231 cells. **C.** Dose-dependent differential effects of Sodium butyrate(SB) and trichostatin A (TSA) on the expression of DNMT3A protein in MDA-MB-231 cells. **D.** Effect of Sodium butyrate (SB) and trichostatin A (TSA) on the expression of H3K27me3 in MCF10A, MDA-MB-231 cells.

To gain the underlying insights regarding the differential effects of SB and TSA on global DNA methylation, we next determined their potential effects on three DNMTs in these cells. The results indicated increased levels of DNMT1 and DNMT3B in cancer cells when compared to normal cells but not DNMT3A levels which were low in cancer cells. Surprisingly, SB induced DNMT3A levels, which might explain DNA hypermethylation (Fig. 3B). On the other hand, TSA reduced all three DNMTs levels, which might be causing global DNA hypomethylation (Fig. 3B). To explore further insights of the differential effects of SB and TSA on DNMT3A, we treated MDA-MB-231 cells with different concentrations of the two drugs. Immunoblot results showed a dose-dependent increase in DNMT3A levels with SB treatments and a dose-dependent decrease in TSA treatment in MDA-MB-231 cells (Fig. 3C). Collectively, interpretation of data presented in Fig 3A-C points to a notion that inhibitor drugs-impacted DNMT3A is key for gained hypomethylation or hypermethylation [29].

With induced global DNA hypermethylation upon SB treatment, we hypothesized the induced formation of heterochromatin of many genomic loci. To test our hypothesis, we examined the level of histone methylation H3 lysine 27 trimethylation (H3K27me3), a repressive histone mark and also hallmark of Xi chromosome [30]. Indeed, we observed the significantly decreased levels of H3K27me3 in MDA-MB-231 cells; however, SB treatment caused significant elevation of H3K27me3. In contrast, the TSA treatment did not show such elevation (Fig. 3D). These data suggest that SB and TSA treatments had distinct effects on chromatin structure, expanding and diminishing, respectively. Our data are also consistent with expected consequence of DNA hypermethylation and hypomethylation (Fig. 3A) on impacting chromatin structure.

### Sodium butyrate induces DNMT3A expression by inhibiting UHRF1

Of late it has been demonstrated that UHRF1 ubiquitin ligase targets DNMT3A for ubiquitination and subsequent degradation thereby leading to hypomethylation of DNA in cancer cells[31]. Herein our data demonstrated SB-induced protein levels of DNMT3A (Fig. 3C); therefore, we analyzed the protein levels of UHRF1. The immunoblot results demonstrated a decrease in UHRF1 protein levels when treated with SB (Fig. 4A). To further explore the effect of SB on UHRF1, we performed a dose-dependent study. The immunoblot showed a significant decrease in UHRF1 protein levels in the presence of increasing concentrations of SB, whereas TSA seemed to increase UHRF1 levels at concentration 0.2μM (Fig. 4B). Collectively, we conclude that SB-induced accumulation of DNMT3A may be resulted from SB-inhibited UHRF1.

**Fig 4:**
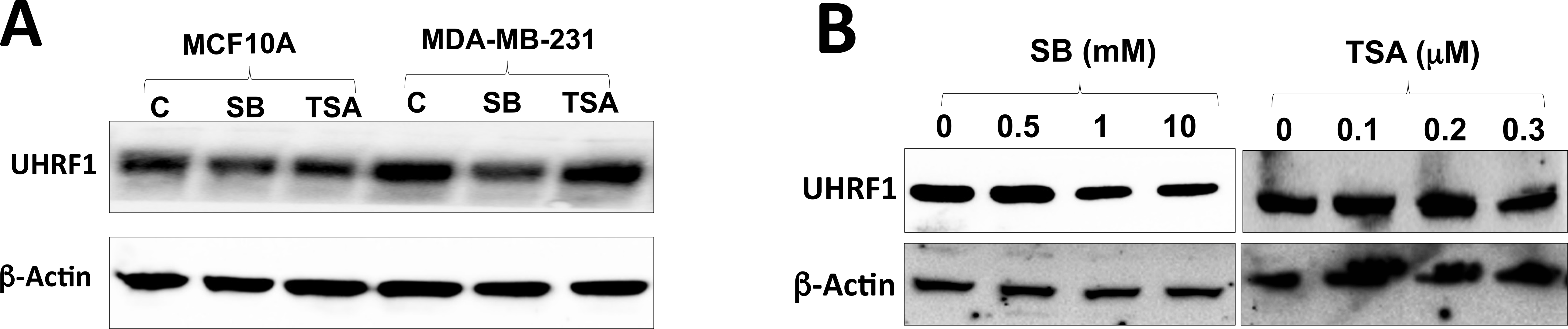
SB induces DNMT3A expression by inhibiting UHRF1 protein. **A.** UHRF1 protein levels inMCF10A, MDA-MB-231 cells in response to Sodium butyrate (SB) and Trichostatin A (TSA). **B.** Immunoblot showing the UHRF1 protein levels in response to different doses of SB and TSA treatment in MDA-MB-231 cells.

### Sodium butyrate reverses the expression of differentially expressed genes

To a deeper insight into the differential effects of SB and TSA on gene expression, we did RNA-seq analyses of MCF10A and MDA-MB-231 cells treated with or without the drugs. Although the data is from a single data set, the heat map clearly indicates differential effects of both drugs in normal and cancerous breast cells (Fig. 5A). Out of 1213 genes that showed differential expression and enrichment in cancer cells (MDA-MB-231) compared to MCF10A normal cells, 502 genes were downregulated and 711 genes were upregulated significantly (Supplementary file 1). Similarly, out of 1052 differentially expressed genes in SB-treated MDA-MB-231 cells, 503 genes were down regulated and 549 were up regulated compared to SB-treated MCF10A cells. In TSA-treated MDA-MB-231 cells, 452 genes were down regulated and 404 were upregulated compared to TSA-treated MCF10A cells. The SB-treated and TSA-treated MDA-MB-231 cells showed very few differentially expressed genes (only six genes that are upregulated in SB-treated cells) and the SB-treated and TSA-treated MCF10A cells showed five down regulated and 33 upregulated genes in SB-treated cells (Supplementary file 1). A detailed analysis of the RNA-seq data indicated that the genes known to be downregulated due to promoter hypermethylation and genes overexpressed due to promoter hypomethylation in in breast cancer, were upregulated in SB-treated MDA-MB-231 cells as compared to untreated cells (Table 3). TSA treatment also showed similar effects on few genes (Table 3); However, SB treatment showed more differential effects.

Data in Fig 2 and 3 linked SB and TSA treatments to different effects on the expression of *XIST* and *TSIX*. We therefore next focused on X-chromosome-specific genes for more insights. RNA-seq analyses showed a marked difference in expression of genes between normal and cancerous breast cells (Fig. 5B). Further analysis of the X-chromosome-specific genes revealed that genes that were upregulated in MDA-MB-231 cells compared to MCF10A cells were downregulated in SB-treated MDA-MB-231 cells and *vice-versa*, indicating SB’s role in re-inactivation of X chromosome (Table 3). The tumor suppressor genes such as PHRF1, WT, SEMA3A and PER3 are differentially methylated upon SB treatment.

**Fig 5:**
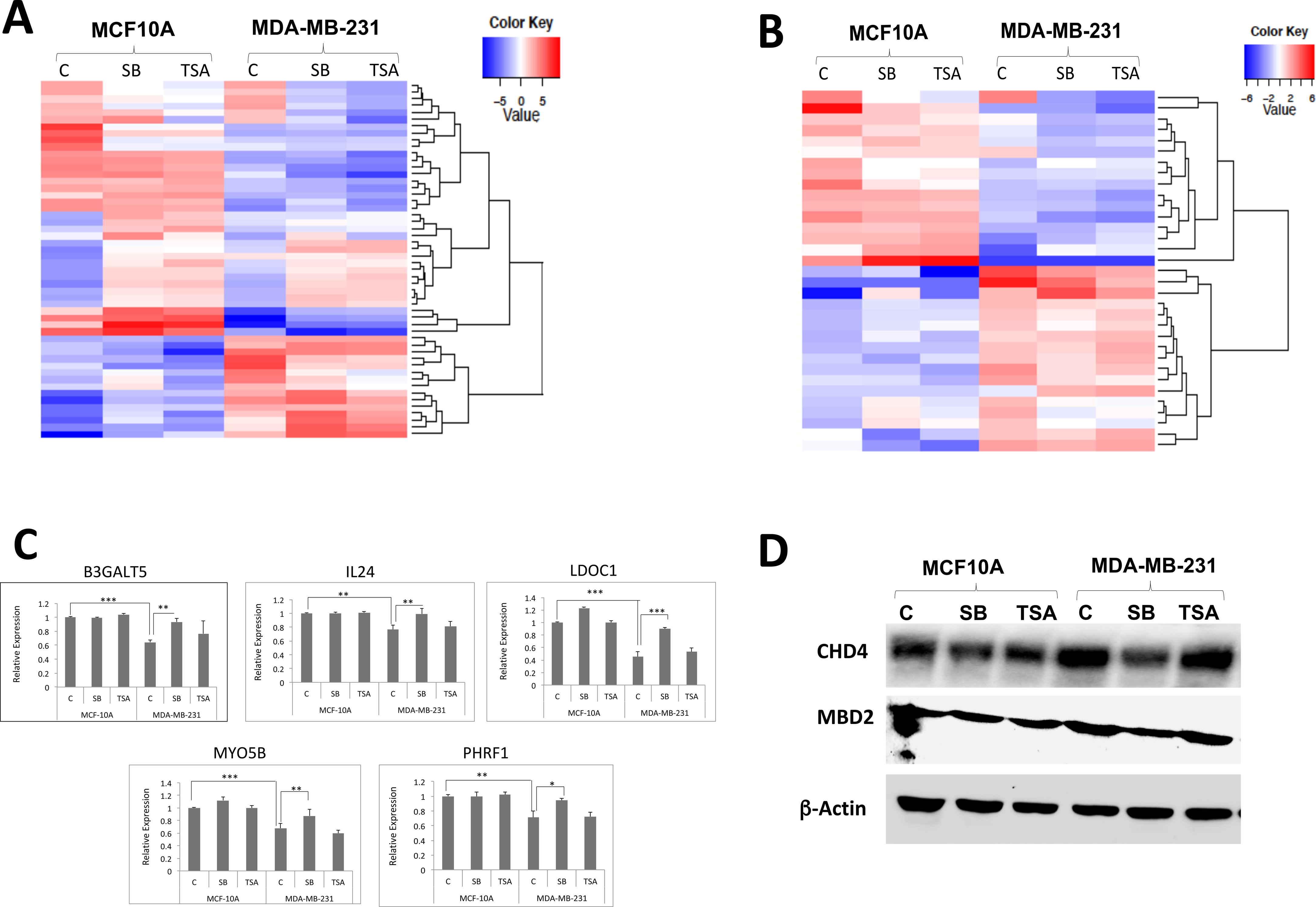
SB reverses the expression of differentially expressed genes. **A.** Effect of Sodium butyrate (SB) and trichostatin A (TSA) on global RNA expression in MCF10A and MDAMB-231 cells. **B.** Heat map showing the differential gene expression of X-chromosome specific genes. A distinct clustering of genes is noted in normal and cancerous cells. **C.** Validation of RNA-seq results for 5 genes by real-time PCR. **D.** Protein levels of subunits of NuRD complex in the nuclear fractions of MCF10A, MDA-MB-231 cells.

The RNA-seq data was further validated by qRT-PCR for some of the genes listed in Table 3 and 4 (Fig. 5C).

Nucleosome remodeling and deacetylase complex (NuRD) consisting of HDAC1, HDAC2, MTA1, MBD2, and CHD4 is known to be involved in chromatin repression and gene silencing. Among complex members, MBD2 is known to repress *XIST* expression *via* DNA hypermethylation [32]. Because SB treatment increased *XIST* expression along with global DNA hypermethylation *via* DNMT3A induction, we analyzed the expression levels of some of the NuRD complex subunits in normal and cancerous breast cells. The immunoblot results demonstrated an overexpression of NuRD subunits in cancer cells and SB down regulated the expression of these proteins (Fig. 5D).

## Discussion

Dosage compensation is a phenomenon known to equalize the gene expression on sex chromosomes, specifically X chromosome in males and females. There are several mechanisms by which dosage compensation is achieved in different organisms [33]. In mammals, dosage compensation is attained by random inactivation of one of the two X chromosomes in females and the process is known as X chromosome inactivation (XCI) [1]. Inactivation is established by painting of the Xi chromosome with XIST long noncoding RNA (lncRNA) in *cis* followed by DNA hypermethylation, enrichment of inactive histone modifications such as H3K27me3 and histone hypoacetylation. Extensive research has established the role of DNMT1 in DNA hypermethylation and localization of XIST onto Xi by BRCA1 thus maintaining the Xi status and dosage compensation. Both the epigenetic mechanisms of DNA hypermethylation and histone hypoacetylation are important to maintain gene silencing and Xi status. The present study aimed at understanding the role of HDACs in Xi maintenance using normal mammary epithelial cells (MCF10A) where there is one Xa and one Xi and cancerous breast cells (MDA-MB-231) where there is skewed Xi status in presence or absence of HDAC inhibitors, SB and TSA.

Of the four classes of HDACs identified in humans, Class I, II and IV are Zinc-dependent enzymes, which are known to be involved in gene regulation *via* histone deacetylation. However, Class III HDACs, Sirtuins, are NAD-dependent enzymes that are mainly involved in metabolic homeostasis and therefore histone deacetylation is not their primary function. Class IV consists of a single member, HDAC11 with unclear function. Class II HDACs shuttle between nucleus and cytoplasm with histone and non-histone cellular proteins as substrates. Class I HDACs therefore are involved only with histone deacetylation, chromatin remodeling and gene regulation. In the present study, first, the expression of Class I and Class II HDACs was studied to assess the differential expression of HDACs upon drug treatments. Then the interested HDACs that might have a role in Xi maintenance in the two cell lines were pursued further. The results showing inhibition of HDAC1, HDAC2, HDAC4 and HDAC6 overexpressed HDACs in cancer cells by SB and TSA clearly indicate the possibility of these HDACs in Xi maintenance. Our results are in agreement with previous studies where valproic acid induced HDAC2 protein degradation in breast cancer cells [34]. Among the four HDACs, HDAC1 and HDAC2 are exclusively in nucleus and are components of nuclear remodeling and deacetylase complex (NuRD) known to be involved in chromatin remodeling [35]. Overexpression of these HDACs gives a clue of their role in Xi chromatin remodeling in cancer cells leading to its reactivation. HDAC6, a class II HDAC, is an X-chromosome-specific and estrogen-regulated gene [36]. Our ChIP-seq results demonstrate HDAC6 important for regulation of transcription elongation of many genes, particularly immediate early genes such as *c-Jun* [*13*]. Previous studies also reported overexpression of HDAC6 in breast cancer is linked to poor survival [37]. In the present study, HDAC6, both mRNA and protein, is overexpressed in MDA-MB-231 cells and the levels of expression seemed to decrease a bit when treated with SB and TSA indicating its expression linked to Xi reactivation.

To understand the role of HDACs in Xi reactivation, we studied the mRNA expression of Xi-specific gene *XIST* and Xa-specific genes *TSIX*, *PGK1* and *G6PD* [38]. Similar to previous studies reporting lack of *XIST* expression in some of the breast cancer cells [39], our results demonstrated no *XIST* or extremely low expression in MDA-MB-231 cells (Fig. 2A). However, the transcript levels of *TSIX* were shown to be much higher in MDA-MB-231 cells compared to MCF10A cells. This indicates that the cancer cells adopt the expression of *TSIX* to inhibit XIST on Xi leading to Xa duplication. *TSIX* expression is significantly decreased upon HDAC inhibition further defining the role of HDACs in Xi maintenance. Similarly other Xa-specific genes, *PGK1* and *G6PD* expression was reduced in SB and TSA-treated cells supporting the hypothesis that HDACs play a role in Xi maintenance.

*BRCA1* is a tumor suppressor gene involved in DNA repair and transcription regulation by interacting with histone deacetylases. Loss or mutation of *BRCA1* leads to breast cancer development[40]. Earlier it was reported that BRCA1 regulates *XIST* RNA expression and thus regulates Xi heterochromatin [22, 41]. Additional studies have shown that triple-negative basal-like breast cancers are more aggressive with loss of *BRCA1* and Xi reactivation [42]. In the present study, we observed decreased expression of *BRCA1* in MDA-MB-231, a basal-like breast cancer cell line with wild type BRCA1 [43]. HDAC inhibition with SB caused increased BRCA1 levels at RNA and protein level in MDA-MB-231 while TSA reduced the levels. H3K56ac levels are recognized by BRCA1 at the site of DNA double strand break. We therefore also analyzed the levels of H3K56ac in response to HDAC inhibitors, which demonstrated an increased level as observed in previous studies [44] [45].

DNA hypermethylation is one of the hallmarks of Xi chromosome and global DNA hypomethylation is a characteristic feature in cancer. We next studied effect of HDAC inhibition on global methylation levels of MCF10A and MDA-MB-231 cells. As known, the cancer cells showed decreased levels of DNA methylation. However, a significant increase in global methylation levels in all the cells when treated with SB was observed. These results are in agreement with previous studies [46]. On the other hand, TSA caused global hypomethylation as known earlier [47]. Recently, it is demonstrated that the number of methylated CpGs on X chromosomes are significantly reduced in breast cancer cells compared to normal breast epithelial cells [48]. Other study reported that the Xa and Xi differ in DNA methylation status specifically at CpG islands [49]. Since DNMTs are responsible for DNA methylation, we also studied the protein expression levels of DNMT1, DNMT3A and DNMT3B in these cell lines. A significant increase in DNMT1 and DNMT3B protein levels was observed in MDA-MB-231 cells when compared to MCF10A cells. These results from our study are in well agreement with previous studies showing similar results [50–52]. The levels of DNMT1 and DNMT3B were decreased when treated with SB and TSA as observed previously [51]. However, the hypermethylation of DNA caused by SB might be due to the induction of DNMT3A. Although the hypermethylation effect of SB is known since long time, the mechanism remained unexplored and we for the first time demonstrate that SB induce DNMT3A, a de novo methyltransferase, which might be causing global DNA hypermethylation.

We also analyzed the H3K27 trimethylation, a characteristic feature of Xi chromosome by Western blot analysis. Breast cancer cell lines showed significant low levels of H3K27me3 levels which increased upon HDACi treatment implicating role of HDACs in restoring DNMTs normal expression and global DNA methylation levels leading to maintenance of Xi. Recently it is reported that UHRF1 is generally overexpressed in cancer cells and that it causes degradation of DNMT3A leading to hypomethylation of DNA [31]. We therefore analysed UHRF1 protein levels and observed that UHRF1 is up regulated in cancer and that SB-treatment significantly down regulated the levels of UHRF1 in cancer cells thus inducing DNMT3A levels.

Aberrant DNA methylation leading to hypermethylation of promoter-enhancer regions of tumor suppressor genes and hypomethylation of genes involved in progression is a well-known phenomenon in breast cancer [53]. We had performed RNA-seq of MCF10A and MDA-MB-231 cells treated with or with SB and TSA to gain an insight into the differential gene expression. The RNA-seq analysis revealed that SB induced the expression of several tumor suppressor genes were repressed due to hypermethylation or Loss of Heterozygosity (LOH) as indicated in Table 3. The X-chromosome methylation pattern also has been affected as indicated by the increased *LDOC1* expression, a gene whose promoter known to be hypermethylated in cancer [54]. However, a detailed methylation-specific sequencing and analysis needs to be carried out to further gain a deeper insight onto differential methylation effects of SB.

In view of the role of NuRD complex in cancer progression and skewed Xi [32, 55], given the functional significance of HDAC1 and HDAC2 in NuRD complex and inhibition of HDAC1 and HDAC2 by sodium butyrate in the present study, we also analyzed NuRD complex subunits (CHD4 and MBD2) by immunoblot. The HDAC inhibitor treatment significantly down regulated these proteins indicating possible involvement of HDAC1 and HDAC2 in Xi maintenance. Inhibition of HDAC1 and/or HDAC2 for the treatment of cancer has been the recent research focus [56].

In conclusion, our data indicates that HDACs, specifically HDAC1 and HDAC2 of Class I HDACs, play an important role in maintenance of Xi. However, in breast cancer cells, overexpression of DNMT1 along with Class I HDACs cause aberrant DNA methylation and chromatin remodeling leading to Xi reactivation and increased malignancy (Fig.6). Inhibition of HDAC1 and HDAC2 specifically by SB restored normal DNA methylation levels by inducing DNMT3A and also degrading HDAC2 protein levels leading to dissolution of NuRD complex and thus modified chromatin structure. On the other hand, TSA, which mostly inhibits HDAC1, HDAC4, and HDAC6, did not show such beneficial effect of Xi restoration. However, further in depth studies are in progress on the tumor suppressor gene promoter methylation levels and CpG methylation levels in these cells in presence and absence of SB and TSA inhibitor in order to confirm the therapeutic potential of given HDAC inhibitor to treat high grade breast tumors.

**Fig 6:**
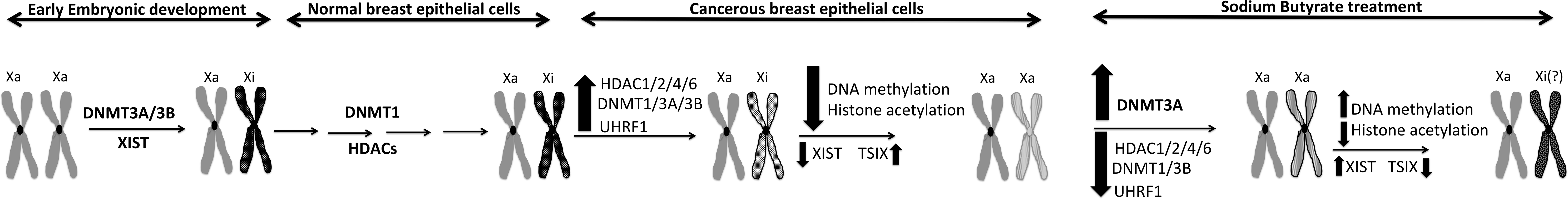
Schematic representation of the probable effects of sodium butyrate in restoring Xi in breast cancer cells.

## Materials and methods

### Cell culture and treatments

Normal human mammary epithelial cells, MCF10A and breast cancer cell line, MDA-MB-231, were generous gift from colleague Dr. Saraswati Sukumar (JHU), who obtained from ATCC recently. MCF10A cells (passage number 6) were cultured in DMEM/F12 medium (Gibco) supplemented with 5% horse serum (Gibco), 5% FBS (Sigma), penicillin streptomycin solution (1x) (Gibco) and mammary epithelial growth supplement (1x) (Gibco) at 37 °C and 5% CO_2_. MDA-MB-231 (passage number 6) were grown in DMEM medium (Gibco) supplemented with 10% FBS (Sigma) and penicillin streptomycin solution (1x) (Gibco) at 37 °C and 5% CO_2_. All cells were sub-cultured twice a week. A stock solution of 1 M of Sodium butyrate (SB) (Sigma) in sterile water and 100 mM Trichostatin A (TSA) (Sigma) in DMSO was prepared and stored at −20 °C. Cells were treated with SB and TSA at a final concentration of 1mM and 100 nM for 24 h.

### Immunoblot

Cells were pelleted by centrifugation at 2000 rpm, 3 min and washed once in ice-cold PBS. Cells were lysed in RIPA buffer (Sigma) consisting of 1x protease inhibitor cocktail (Sigma) and 1x phosphatase inhibitor cocktail (Sigma) for 30 min on ice. The total cell lysate was isolated by centrifugation at 13000 rpm, 4 °C,20 min and Qubit estimated the total protein concentration according to manufacturer’s protocol. A total of 50 μg of protein was separated on 4-15% SDS gel (Bio-Rad), transferred onto PVDF membrane and probed with primary antibody (Table 1) at a dilution of 1:1000 prepared in 5% blotting grade blocker (Bio-Rad) for 12-16 h at 4 °C. The protein bands were visualized after secondary antibody incubation (1:10000) for 1h at room temperature followed by ECL detection (GE Amersham).

**Table 1:** List of antibodies used in the study

| **Antibody** | **Catalog #** | **Source** |
| --- | --- | --- |
| HDAC1 | 06-720 | Upstate |
| HDAC2 | 07-222 | Upstate |
| HDAC3 | sc-11417 | Santa Cruz |
| HDAC4 | sc-11418 | Santa Cruz |
| HDAC5 | sc-11419 | Santa Cruz |
| HDAC6 | sc-11420 | Santa Cruz |
| HDAC8 | sc-11405 | Santa Cruz |
| HDAC10 | sc-54215 | Santa Cruz |
| β-Actin | A5441 | Sigma |
| BRCA1 | 9010P | Cell Signaling |
| DNMT1 | A300-042 | Bethyl |
| DNMT3A | Ab2850 | Abcam |
| DNMT3B | IMG-184A | Imginex |
| 5-mC | A-1014-050 | Epigentek |
| CHD4 | A301-083A | Bethyl |
| UHRF1 | AP8846c | Abgent |
| H3K9ac | Ab4441 | Abcam |
| MTA2 | A300-395A | Bethyl |
| H3K56ac | 07-677 | Upstate |
| Histone H3 | Ab1791 | Abcam |
| MBD2 | 07-198 | Upstate |

### RNA isolation and RT-PCR

Total RNA was isolated from cells using Trizol reagent following manufacturer’s protocol. The concentration of total RNA was estimated by Qubit and 1 μg of RNA was reverse transcribed into cDNA using Superscript III First strand cDNA synthesis kit (Invitrogen). 2 μl of out of 20 μl obtained cDNA was used for PCR using gene specific primers (Table 2).

**Table 2:**
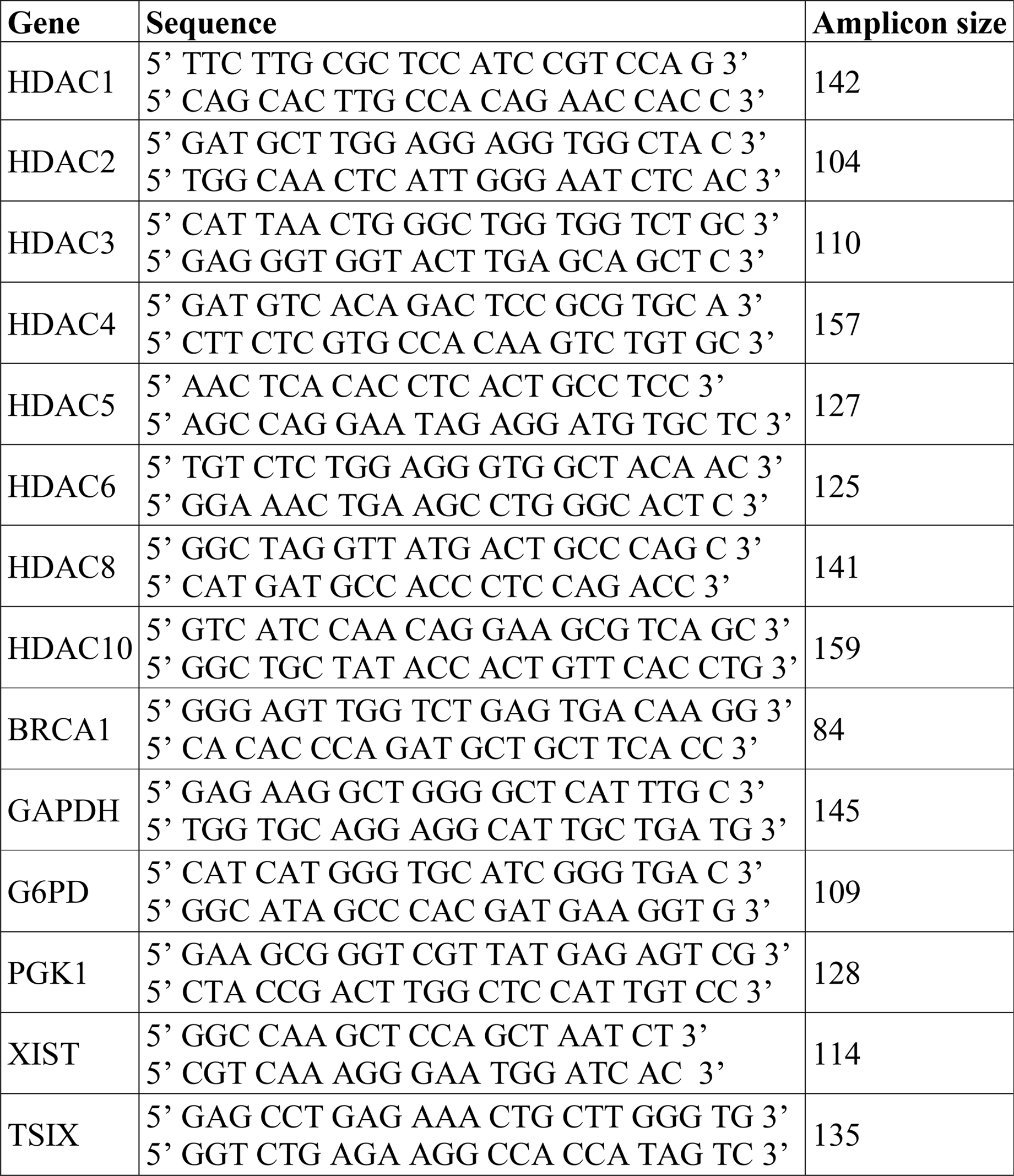
Primer sequences used in the study for RT-PCR analysis

**Table 3:**
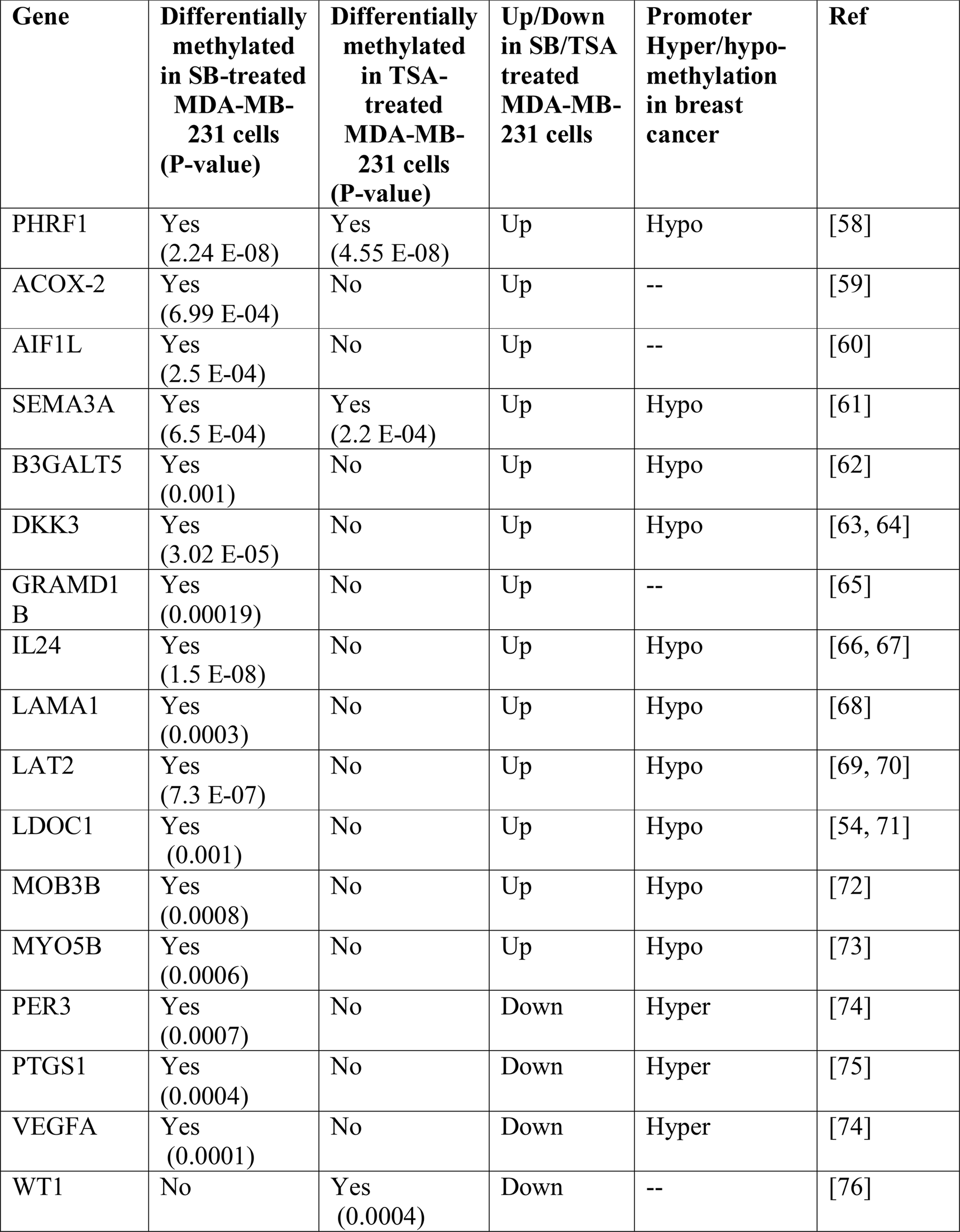
Partial list of differentially expressed Tumor suppressor genes in SB-treated MDA-MB-231 cells compared to MCF10A

| **Gene** | **Differentially methylated in SB-treated MDA-MB-231 cells (P-value)** | **Differentially methylated in TSA-treated MDA-MB-231 cells (P-value)** | **Up/Down in SB/TSA treated MDA-MB-231 cells** | **Promoter Hyper/hypo-methylation in breast cancer** | **Ref** |
| --- | --- | --- | --- | --- | --- |
| PHRF1 | Yes (2.24 E-08) | Yes (4.55 E-08) | Up | Hypo | [58] |
| ACOX-2 | Yes (6.99 E-04) | No | Up | -- | [59] |
| AIF1L | Yes (2.5 E-04) | No | Up | -- | [60] |
| SEMA3A | Yes (6.5 E-04) | Yes (2.2 E-04) | Up | Hypo | [61] |
| B3GALT5 | Yes (0.001) | No | Up | Hypo | [62] |
| DKK3 | Yes (3.02 E-05) | No | Up | Hypo | [63, 64] |
| GRAMD1 B | Yes (0.00019) | No | Up | -- | [65] |
| IL24 | Yes (1.5 E-08) | No | Up | Hypo | [66, 67] |
| LAMA1 | Yes (0.0003) | No | Up | Hypo | [68] |
| LAT2 | Yes (7.3 E-07) | No | Up | Hypo | [69, 70] |
| LDOC1 | Yes (0.001) | No | Up | Hypo | [54, 71] |
| MOB3B | Yes (0.0008) | No | Up | Hypo | [72] |
| MYO5B | Yes (0.0006) | No | Up | Hypo | [73] |
| PER3 | Yes (0.0007) | No | Down | Hyper | [74] |
| PTGS1 | Yes (0.0004) | No | Down | Hyper | [75] |
| VEGFA | Yes (0.0001) | No | Down | Hyper | [74] |
| WT1 | No | Yes (0.0004) | Down | -- | [76] |

### Dot blot

Genomic DNA was isolated from cells and a dot blot was carried out using 60 ng and 30 ng DNA and anti-5mC antibody as described previously [57].

### RNA-seq by Illumina HiSeq

Total RNA of each sample was extracted using TRIzol Reagent (Invitrogen)/RNeasy Mini Kit (Qiagen)/other kits and was quantified and qualified by Agilent 2100 Bioanalyzer (Agilent Technologies, Palo Alto, CA, USA), NanoDrop (Thermo Fisher Scientific Inc.) and 1% agrose gel. 1 μg total RNA with RIN value above 7 was used for library preparation. Next generation sequencing libraries were constructed according to the manufacturer’s protocol (NEBNext^®^ Ultra^TM^ RNA Library Prep Kit for Illumina^®^). The poly(A) mRNA isolation was performed using NEBNext Poly(A) mRNA Magnetic Isolation Module (NEB) or Ribo-Zero^TM^ rRNA removal Kit (illumina). The mRNA fragmentation and priming was performed using NEBNext First Strand Synthesis Reaction Buffer and NEBNext Random Primers. First strand cDNA was synthesized using ProtoScript II Reverse Transcriptase and the second-strand cDNA was synthesized using Second Strand Synthesis Enzyme Mix. The purified (by AxyPrep Mag PCR Clean-up (Axygen)) double-stranded cDNA was then treated with End Prep Enzyme Mix to repair both ends and add a dA-tailing in one reaction, followed by a T-A ligation to add adaptors to both ends. Size selection of Adaptor-ligated DNA was then performed using AxyPrep Mag PCR Clean-up (Axygen), and fragments of ~360 bp (with the approximate insert size of 300 bp) were recovered. Each sample was then amplified by PCR for 11 cycles using P5 and P7 primers, with both primers carrying sequences which can anneal with flow cell to perform bridge PCR and P7 primer carrying a six-base index allowing for multiplexing. The PCR products were cleaned up using AxyPrep Mag PCR Clean-up (Axygen), validated using an Agilent 2100 Bioanalyzer (Agilent Technologies, Palo Alto, CA, USA), and quantified by Qubit 2.0 Fluorometer (Invitrogen, Carlsbad, CA, USA). The libraries with different indices were multiplexed and loaded on an Illumina HiSeq instrument according to manufacturer’s instructions (Illumina, San Diego, CA, USA). Sequencing was carried out using a 2×150bp paired-end (PE) configuration; image analysis and base calling were conducted by the HiSeq Control Software (HCS) + RTA 2.7 (Illumina) on the HiSeq instrument. The sequences were processed and analyzed by Nucleome Informatics.

## Acknowledgements

Acknowledge Mr. Bipin Kumar, Nucleome Informatics for RNA-seq.

